# Regulating ferredoxin electron transfer using nanobody and antigen interactions

**DOI:** 10.1101/2024.10.23.619829

**Authors:** Albert Truong, Jonathan J. Silberg

## Abstract

Fission and fusion can be used to generate new regulatory functions in proteins. This approach has been used to create ferredoxins (Fd) whose cellular electron transfer is dependent upon small molecule binding. To investigate whether Fd fragments can be used to monitor macromolecular binding reactions, we investigated the effects of fusing fragments of *Mastigocladus laminosus* Fd to nanobodies and their protein antigens. When Fd fragments arising from fission were fused to green fluorescent protein (GFP) and three different anti-GFP nanobodies, split proteins were identified that supported Fd-mediated electron transfer from Fd-NADP reductase (FNR) to sulfite reductase (SIR) in *Escherichia coli*. However, the order of nanobody and antigen fusion to the Fd fragments affected cellular electron transfer. Insertion of these anti-GFP nanobodies within Fd had differing effects on electron transfer. One domain-insertion variant was unable to support cellular electron transfer unless it was coexpressed with GFP, while others supported electron transfer in the absence of GFP. These findings show how Fds can be engineered so that their electron transfer is regulated by macromolecules, and they reveal the importance of exploring different nanobody homologs and fusion strategies when engineering biomolecular switches.

## Introduction

Iron-sulfur (Fe-S) cluster containing ferredoxins (Fds) are abundant across the tree of life, with individual species frequently containing multiple paralogs.^1,2^ Fds have evolved to support electron transfer (ET) between biochemical pathways important for energy transduction, such as hydrogen and alcohol production, carbon and nitrogen fixation, and sulfur assimilation.^3,4^ These low potential iron-sulfur proteins are thought to function as central energy-conserving ET hubs, distributing electrons to various metabolic pathways through protein-protein interactions with over a hundred classes of partner proteins.^5–8^ Biophysical studies have revealed that these oxidoreductases evolved sequences that fold into a variety of structures and coordinate either 2Fe-2S, 3Fe-4S, or 4Fe-4S clusters,^3,4^ and electrochemical studies have shown that Fds present a range of midpoint potentials.^9–11^ In addition, biochemical studies have found that Fds present a range of efficiencies when coupling with different partner proteins.^12–18^ While Fds play a central role in controlling energy flow across the electron fluxome, we cannot yet easily control the proportion of electrons relayed by Fds among different pathways in cells.

The large array of Fd partners and ability to transfer low potential electrons has led to their use in a variety of synthetic biology applications, including metabolic engineering,^19–21^ light-driven biosynthesis,^22–25^ optogenetics,^26^ fluorescent reporting,^27^ and bioelectronic reporting.^28^ Fds have also been targeted for protein engineering to test theories about their evolution.^29^ *In vitro* studies have shown that small iron-sulfur cluster-containing peptides can be synthesized and used to support multiple oxidation−reduction cycles.^30,31^ In addition, iron-sulfur binding sites have been incorporated into synthetic coiled-coil structures,^32^ and these proteins have been shown to acquire iron-sulfur clusters in cells. Further, Fds have been designed to mimic ancestral Fds arising from a peptide-duplication event.^29^ When expressed in bacteria, these computationally-designed Fds can provide the reductive power needed for nutrient assimilation.^33^ Chimeric Fds have also been created to understand how structure controls midpoint potential and thermostability.^34,35^ Finally, Fds have been modified to create protein switches that regulate cellular ET for fast sensing applications. These efforts have identified numerous Fd designs that present ligand-dependent ET in cells.^36,37^ To date, these protein design efforts have only generated Fd switches that respond to small molecule analytes.

One strategy used to engineer proteins whose activities are controlled by macromolecular binding is to generate single domain antibodies, referred to as nanobodies (Nbs),^38^ and to fuse those Nbs to other proteins. Through this process, protein activities have been regulated, such as luminescence^39,40^ and transcription.^41,42^ Central to the success of these design efforts is understanding how to fuse Nbs in a way that makes the activity of the modified protein conditional upon antigen binding. To better understand how such regulation can be accomplished with a protein electron carrier, we investigated how three different anti-GFP Nbs can be fused to Fd fragments to regulate cellular ET. We identify Nb and GFP fusions strategies that support Fd-fragment complementation. We also discover a Nb-inserted Fd whose ET can be switched on by GFP expression in *Escherichia coli*.

## Experimental Section

### Materials

Chemicals for growth medium were from VWR, MilliporeSigma, Fisher, Apex Biosciences, Research Products International, or BD Biosciences. Chloramphenicol, streptomycin sulfate, kanamycin, and isopropyl β-D-1-thiogalactopyranoside (IPTG) were from Research Products International. Anhydrotetracycline (aTc) and 3-oxododecanoyl-L-homoserine lactone (AHL) were from Sigma-Aldrich. Enzymes and molecular biology kits were from Zymo Research, Qiagen, and New England Biolabs.

### Plasmid construction

All plasmids are listed in Table S1. Plasmids encoding *Mastigocladus laminosus* Fd (pFd007), a C42A Fd mutant (pFd007), a split Fd fused to FKBP/FRB (pRAP007.35), and *Zea mays* FNR and SIR (pSAC01) were previously described.^37^ A plasmid for AHL-inducible expression of GFP (pSH001) was generated, which uses LasR to regulate GFP expression.^43^ Plasmids for expressing Fds split after residue 35 were made by replacing the FKBP and FRB coding sequences in pRAP007.35 with GFP and the anti-GFP Nbs LaG-2, (pAT020 and pAT021), LaG-16 (pAT022 and pAT023), and LaG-41 (pAT024 and pAT025).^38^ The genes encoding the Nbs were synthesized commercially by Integrated DNA Technologies, while the gene encoding GFP was PCR amplified from pSH001. Plasmids for expressing Fds split after residue 55 (pAT026 to pAT0031) were made by amplifying the DNA inserted into the Fd gene from pAT020 to pAT025 and cloning these amplicons after the codon encoding Fd residue 55 in pFd007. Vectors for expressing split Fds with different linker lengths (pAT032 to pAT039) were created by linearizing pAT020 and pAT021 through PCR amplification using primers that altered the linker sequences, and the amplicons were circularized using Golden Gate cloning.^44^ Vectors for expressing Fds having Nb domains inserted (pAT040, pAT041, and pAT042) were made by PCR amplifying pAT020, pAT022, and pAT024 in a way that deletes the GFP and regulatory sequences and circularizing the amplicons. A vector for inducible GFP expression (pAT019) was made that is compatible with pAT040, pAT041, and pAT042 by cloning a pSC101 origin in place of ColE1 origin. All plasmids were sequence verified using Sanger sequencing (Azenta Life Sciences).

### Growth medium and strains

*E. coli* XL1-Blue (Agilent, Inc.) was used for all cloning, while *E. coli* EW11 was used assay Fd electron transfer.^45^ For molecular cloning, cells were grown in lysogeny broth (LB). For the complementation studies, M9 complete (M9c) and M9 selective (M9sa) media were made as described.^46^ *E. coli* EW11 can grow in the absence of Fd in M9c, but it cannot grow in M9sa in the absence of a Fd that mediates ET from Fd-NADP reductase (FNR) and sulfite reductase (SIR).^46^

### Growth complementation

To measure growth complementation of *E. coli* EW11, cells were transformed using electroporation with plasmids expressing different Fds and pSAC01, a plasmid that constitutively expresses *Zea mays* FNR and SIR.^37^ With plasmids expressing split Fds, protein fragments were expressed using aTc- and IPTG-inducible promoters. Cells were grown on LB-agar plates containing 34 ug/mL chloramphenicol and 100 ug/mL streptomycin. Individual colonies were then picked into M9c (1 mL) containing antibiotics (34 ug/mL chloramphenicol and 100 ug/mL streptomycin) and the indicated inducer concentrations. After growing cultures overnight at 37°C while shaking at 250 rpm, cultures were diluted 1:500 into M9sa medium (1 mL) containing antibiotics and inducers. These cultures were then grown at 37°C under similar shaking conditions for 96 hours. Every 24 hours, an aliquot (100 µL) was removed measure optical density (absorbance at 600 nm) and green fluorescence (excitation: 488 nm, emission: 507 nm) using shallow 96-well plates with a TECAN Spark.

### GFP-regulated electron transfer

To identify optimal GFP induction conditions, *E. coli* EW11 transformed with pAT019 was grown in M9c (500 µL) containing kanamycin (50 ug/mL) and 3-oxo-C12-homoserine lactone (0, 0.1, 1, 10, 100, and 1000 nM). After 20 hours at 37°C, with shaking at 600 rpm, OD and green fluorescence was measured. To assess the effect of GFP expression on the activity of Fds with nanobody insertions, *E. coli* EW11 was transformed with three plasmids, including a plasmid that expresses Fds with Nb insertions (pAT040, pAT041, or pAT042), a plasmid that expresses GFP (pAT019), and a plasmid that expresses FNR and SIR (pSAC01). Cells were grown on LB-agar medium containing 34 ug/mL chloramphenicol, 100 ug/mL streptomycin, and 50 ug/mL kanamycin. Individual colonies were used to inoculate M9c cultures (0.5 mL) containing antibiotics (17 ug/mL chloramphenicol, 50 ug/mL streptomycin, and 25 ug/mL kanamycin) and varying concentrations of aTc (0, 25, 50, and 100 ng/mL) and AHL (0, 1, 10, 100 nM). Cultures were grown for 4.5 hours at 37°C while shaking at 600 rpm. These cultures were diluted 1:167 into M9sa medium (0.5 mL) containing the same antibiotics and inducers. Cultures were grown at 37°C for 48 hours, with similar shaking, and every 24 hours, absorbance and fluorescence were measured.

### Statistics

In all experiments, three or more biological replicates were measured. Growth complementation data is plotted as the mean and standard deviation. To assess significance of growth complementation, a two-tailed Welch’s t-test was used.

## Results & discussion

### Fd-fragment complementation

In a prior study, fragments of *M. laminosus* Fd were identified that are unable to support ET from *Z. mays* FNR to SIR unless the Fd fragments are fused to proteins that assist with fragment complementation.^37^ In those studies, a Fd split after residue 35 (sFd-35) was unable to support cellular ET unless it was fused to peptides that associate into a coiled coil structure, SYNZIP-17 and SYNZIP-18,^47^ and proteins whose association is stabilized by rapamycin, FKBP12 and the FKBP-rapamycin binding domain of mTOR.^48^ To test if Nb and antigen interactions can assist with Fd-fragment complementation, we created constructs for expressing sFd-35 fragments as fusions to GFP and Nbs that bind to different GFP epitopes, including LaG-2, LaG-16 and LaG-41.^38^ The three Nbs were fused to the C-terminus of a Fd fragment containing residues 1 to 35, designated Fd-f1, and GFP was fused to the N-terminus of a Fd fragment containing residues 36 to 99, designated Fd-f2 (Figure 1a). A flexible peptide linker having the sequence (GGGGS)_2_AAA was used to fuse Fd-f1 to GFP and Nbs to Fd-f2. In all constructs, Fd-f1 expression was regulated using an aTc-inducible promoter, while Fd-f2 expression was regulated by an IPTG-inducible promoter.

**Figure 1.**
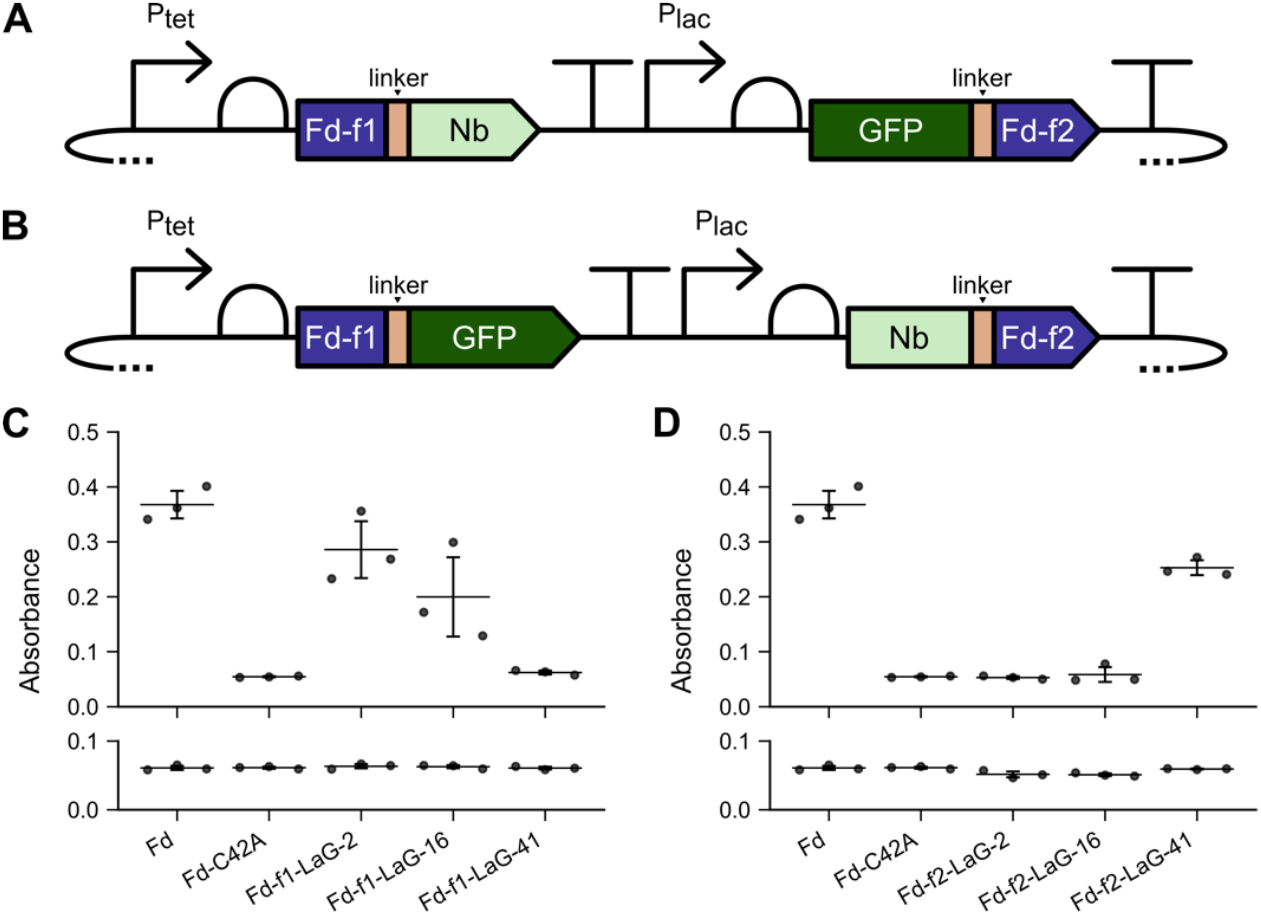
Nb-antigen interactions can assist with Fd-fragment complementation. Protein fragments Fd-f1 and Fd-f2 were created by subjecting the *M. laminosus* Fd to fission after residue 35. The genes encoding these fragments were then fused to the genes encoding anti-GFP Nbs and GFP in two orientations. (**A**) With the first orientation, Fd-f1 was expressed as a fusion to the different Nbs, while Fd-f2 was expressed a fusion to GFP. (**B**) With the second orientation, Fd-f1 was expressed as a fusion to GFP, while Fd-f2 was expressed a fusion to different Nbs. All constructs used *P*_*tet*_ and *P*_*las*_ to regulate the expression of Fd-f1 and Fd-f2, respectively. (**C**) Fd-f1-Nbs and GFP-Fd-f2 and (**D**) Fd-f1-GFP and Nbs-Fd-f2 were expressed in *E. coli* EW11 and growth complementation was measured after 48 hours at 37°C in selective medium containing (*top*) or lacking (*bottom*) the inducers aTc (100 ng/mL) and IPTG (100 nM). As controls, *M. laminosus* Fd and a C42A Fd mutant were expressed using *P*_*tet*_ and assayed for growth complementation in the presence and absence of 100 ng/mL aTc. Data represent the mean and standard deviation of three biological replicates. In panel C, Fd-f1-LaG-2 complements growth significantly more than C42A (p ≤ 0.02; two-tailed Welch’s t-test). In panel D, Fd-f2-LaG-41 complements growth significantly more than C42A (p ≤ 0.002).

To assay whether Nb-antigen interactions can support sFd-35 fragment complementation, each split protein was expressed in *E. coli* EW11.^45^ We found that *M. laminosus* Fd complements growth on selective medium containing sulfate as the only sulfur source after 48 hours, while a mutant Fd (C42A) that cannot coordinate a 2Fe-2S cluster does not complement growth (Figures 1c). In addition, sFd-35 complemented *E. coli* EW11 growth in selective medium when the N-terminal fragment was fused to Nbs LaG-2 and LaG-16, and the C-terminal fragment was fused to GFP. In contrast, the Nb LaG-41 and GFP could not support sFd-35 ET when fused in a similar way. When cells were incubated for longer durations (Figure S1a), the optical density of cells expressing the LaG-2 and LaG-16 designs increased, while there was no change in the cells expressing the LaG-41 design. In all cases, growth complementation was only observed in the presence of the inducers aTc and IPTG. When growth was assayed on non-selective growth medium, strains expressing all designs grew to similar extents in the presence and absence of the inducers (Figure S2). These findings show that Nbs vary in their ability to support sFd-35 fragment complementation when the anti-GFP Nb is fused to Fd-f1 and the GFP antigen is fused to Fd-f2.

To investigate if the fusion orientation affects sFd-35 fragment complementation, an alternative split protein design was studied. In the second design, GFP was fused to the C-terminus of Fd-f1, while the Nbs were fused to the N-terminus of Fd-f2 (Figure 1b). After 48 hours, growth complementation was observed with sFd-35 fragments when fused to LaG-41 and GFP (Figures 1d). This complementation required the addition of the inducers aTc and IPTG. In contrast, growth complementation was not observed after 48 hours when sFd-35 fragments were fused LaG-2 and LaG-16 in either the presence or absence of inducers. After 96 hours, an increase in growth complementation by the LaG-16 design was observed in the presence of inducers (Figure S1b). In non-selective medium, no growth defects were observed (Figure S3). Thus, the LaG-41 design presents the strongest protein-fragment complementation when the Nbs are fused to Fd-f2 and the antigen is fused to Fd-f1. This can be contrasted with sFd-35 designs having the Nbs fused to Fd-f1, where LaG-2 and LaG16 exhibited the strongest complementation.

### Varying the fission site and linkers

To determine if the trends observed arise because of the Fd fragments used for protein design, we targeted a second Fd backbone location for fission. For this design, we subjected *M. laminosus* Fd to backbone fission after residue 55 to create sFd-55. This site was chosen because a prior combinatorial design study found that this split Fd presents strong growth complementation when fused to the termini of a ligand-binding domain.^36^ As with sFd-35, two types of designs were created. First, constructs were created for expressing Nbs as fusions to the C-terminus of Fd-f1 (residues 1 to 55), and GFP was fused to the N-terminus of Fd-f2 (residues 56 to 99). With these split proteins, *E. coli* EW11 growth was only complemented after 24 hours in selective medium when Fd-f1 was fused to Nbs LaG-2 and LaG-16, and Fd-f2 was fused to GFP. Growth complementation was not observed when sFd-55 fragments were fused to LaG-41 and GFP. As with sFd-35, sFd-55 was fused to each Nb and GFP in a second orientation (Figure 2a). With these designs, growth complementation was strongest after 24 hours when the sFd-55 fragments were fused to LaG-41 (Figure 2b). In contrast, growth complementation was not observed in selective medium with the constructs containing LaG2 and LaG-16. With all sFd-55 designs, growth complementation was dependent upon the presence of inducers. Also, all cells grew to a similar density in non-selective medium (Figure S4-S5). These growth complementation trends are identical to those observed with sFd-35, illustrating how the fusion strategy determines which anti-GFP Nbs can support Fd-fragment complementation.

**Figure 2.**
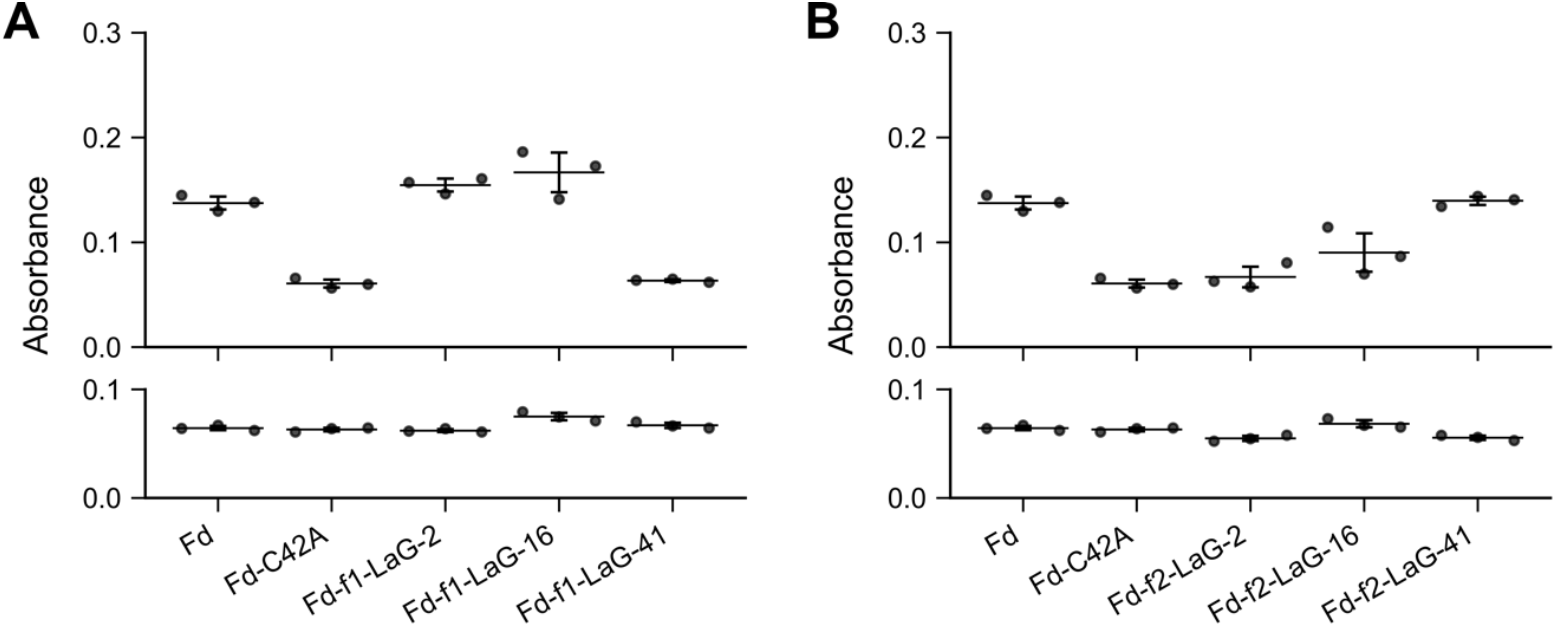
Nb-antigen interactions can be used with a second split Fd to support cellular ET. Fd-f1 and Fd-f2 created by subjecting *M. laminosus* Fd to fission after residue 55 were fused to anti-GFP Nbs and GFP in two orientations as in Figures 1A and 1B. All constructs used P_tet_ and P_las_ to regulate the expression of Fd-f1 and Fd-f2, respectively. (**A**) Fd-f1-Nb and GFP-Fd-f2 and (**B**) Fd-f1-GFP and Nb-Fd-f2 were expressed in *E. coli* EW11 and complementation was measured after 48 hours at 37°C in selective medium containing (*top*) or lacking (*bottom*) the inducers aTc (100 ng/mL) and IPTG (100 nM). As controls, *M. laminosus* Fd and a C42A Fd mutant were expressed using P_tet_ and assayed for growth complementation in the presence or absence of 100 ng/mL aTc. Data represent the mean and standard deviation of three biological replicates. In panel A, Fd-f1-LaG-2 (p < 0.0002) and Fd-f1-LaG-16 (p ≤ 0.01) complement growth significantly more than C42A (two-tailed Welch’s t-test). In panel B, Fd-f2-LaG-41 complements growth significantly more that C42A (p < 0.0001).

In our initial sFd designs, identical linkers were used to fuse Fd-f1 and Fd-f2 to the Nbs and GFP, respectively. This linker had the sequence (GGGGS)_2_AAA. Because prior studies have found that linker design can affect the stability and folding of proteins,^49,50^ we investigated whether changes in linker length alter growth complementation by the sFd-35 fragments having LaG-2 and GFP fused in both orientations. For these constructs, we tested linkers that had been shortened by five residues, having the sequence (GGGGS)_1_AAA, and linkers that had been extended by five residues, with the sequence (GGGGS)_3_AAA. With the designs having the Nbs fused to the C-terminus of Fd-f1 and GFP fused to the N-terminus of Fd-f2, every design complemented the growth of *E. coli* EW11 after 24 hours (Figure 3a). In contrast, the designs having Nbs and GFP fused in the second orientation were unable to complement cell growth on selective medium after similar incubations (Figure 3b). These results show that Fd-fragment complementation does not vary when using linkers that vary in length from eight to eighteen residues.

**Figure 3.**
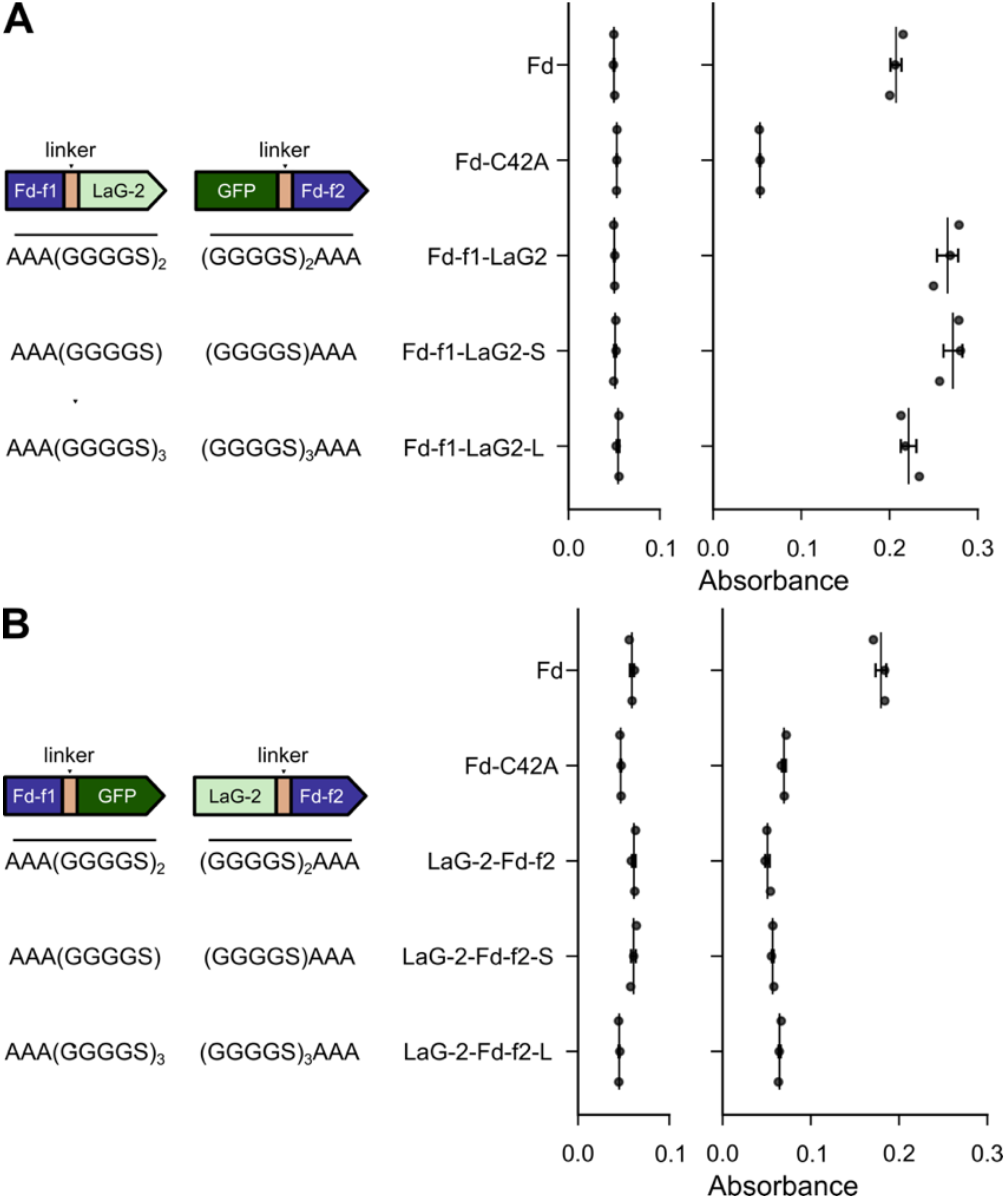
Nb-antigen interactions support Fd-fragment complementation across a range of linker lengths. Growth complementation of *E. coli* EW11 by split Fd designs having (**A**) Fd-f1 fused to LaG-2 and GFP fused to Fd-f2 and (**B**) Fd-f1 fused to GFP and LaG-2 fused to Fd-f2. For each split protein, the linkers used for Fd-f1 (*left*) and Fd-f2 (*right*) are shown. With all constructs, growth complementation was assayed in selective medium containing 100 ng/mL aTc and 100 nM IPTG. Growth complementation was assayed after incubation at 37°C for 24 hours. As controls, *M. laminosus* Fd and Fd C42A were expressed using P_tet_ and assayed for growth complementation in the presence and absence of 100 ng/mL aTc. Data represent the mean and standard deviation of three biological replicates. In panel A, designs having the three different linkers complemented growth significantly more than C42A (p < 0.002; two-tailed Welch’s t-test). In panel B, the average values for each variant were slightly lower than the C42A mutant.

### Nanobody-domain insertion

Domain insertion has been used to create Fd switches whose cellular ET is regulated by endocrine disruptors.^36,37^ These ligand-inducible switches were created by inserting the estrogen receptor ligand-binding domain after residue 35 in *M. laminosus* Fd, which undergoes a large conformational change upon binding estrogen antagonists.^51^ To explore whether Nb insertion at the same site can be used to create Fds whose ET is regulated by a macromolecule, we created constructs for expressing this Fd with LaG-2, LaG-16, and LaG-41 inserted after residue 35 and examined the ability of these proteins to support ET in the absence and presence of GFP. For these experiments, GFP was expressed using an AHL-inducible promoter (Figure S6), while Fds were expressed using an aTc-inducible promoter (Figure 4a).

**Figure 4.**
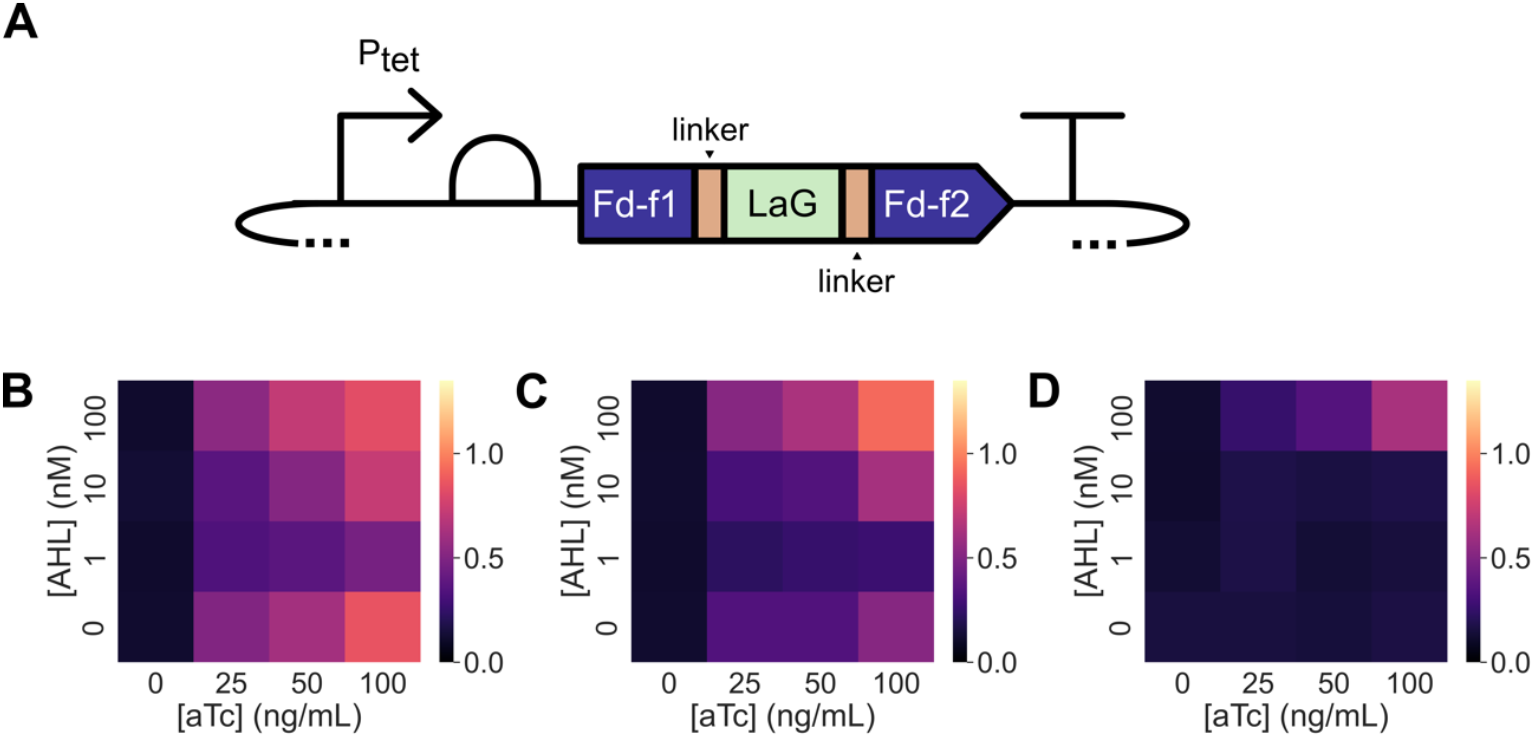
GFP can be used to regulate the ET mediated by a Nb-inserted Fd. (**A**) The Nbs LaG-2, LaG-16, and LaG-41 were inserted into *M. laminosus* Fd after residue 35 to create three different Fd-Nb domain insertion variants. The expression of each Fd-Nb variant was regulated using P_tet_, while GFP expression was regulated using P_las_. Growth complementation was assessed for *E. coli* EW11 containing expression vectors for (**B**) Fd-LaG-2, (**C**) Fd-LaG-16, and (**D**) Fd-LaG-41. Cells were incubated at 37°C for 48 hours in selective medium containing varying concentrations of aTc (0, 25, 50, and 100 ng/mL) and AHL (0, 1, 10, and 100 nM). Data represent the mean of three biological replicates. With Fd-LaG-2 and Fd-LaG-16, in the absence of AHL, a significant increase in growth complementation was induced by aTc (two-tailed Welch’s t-test, p < 0.05) compared with the no aTc condition; aTc did not significantly alter Fd-LaG-41 complementation (p = 0.41) in the absence of AHL. When Fd-LaG-41 expression was induced with 50 ng/mL aTc, 100 µM AHL led to a significant increase in growth complementation compared cells grown in the absence of AHL (two-tailed Welch’s t-test, p < 0.05).

Fd complementation of *E. coli* EW11 growth was first examined in the presence of different GFP-induction conditions. When cells were grown for 48 hours in the presence of a range of aTc concentrations (Figure S7), Fd complemented *E. coli* EW11 growth in selective growth medium to similar extents. Co-induction of GFP expression using AHL had no effect on Fd growth complementation. When similar experiments were performed using Fd variants having LaG-2 and LaG-16 inserted after residue 35, designated Fd-LaG-2 and Fd-LaG-16, growth complementation was observed when protein expression was induced with aTc alone (Figures 4b-c). In contrast, the Fd variant having LaG-41 inserted, Fd-LaG-41, did not complement *E. coli* EW11 growth at any aTc concentration in the absence of AHL (Figure 4d). When experiments were performed in the presence of both aTc and AHL, Fd-LaG-41 complemented cell growth when 100 nM AHL was added to the growth medium. With Fd-LaG-2 and Fd-LaG-16, addition of 1 and 10 nM AHL decreased growth complementation, while 100 nM AHL increased growth complementation. Growth in non-selective medium was similar across all samples (Figure S8). These results show that insertion of the Nb LaG-41 abolishes Fd ET following insertion, while the other Nbs have more modest effects on Fd-mediated growth complementation. Also, these findings show that Fd-LaG-41 ET can be induced in cells by expressing the GFP antigen for the anti-GFP Nb.

## Conclusions

This study shows for the first time that a low potential Fd can be engineered to present cellular ET that is dependent upon expression of another protein. Given that Nbs can be created that bind to a wide range of molecules,^52^ this finding suggests that the domain insertion location targeted in *M. laminosus* can be used for fusion to other Nbs to diversify the molecular recognition that is used to regulate Fd ET. In the future, it will be interesting to explore whether the stability of the Nbs and the Fds targeted for protein design affect the switching activities observed with Fd-Nb fusion proteins. Protein thermostability has been found to affect protein tolerance to topological mutations like fission and permutation,^53–55^ suggesting both Nb and Fd thermostability could affect the function of the Fd-Nb switches and their activities upon binding antigens. Finally, the approach described for regulating cellular ET herein could be used to diversify components for creating living electronics.^56^ To create components for fast electron sensing aqpplications, the protein design approaches described herein could be applied to other classes of protein electron carriers, such as flavodoxins and cytochromes.^3,46,57^

## Author Contributions

Conceptualization, A.T. and J.J.S.; methodology, A.T. and J.J.S.; investigation, A.T.; writing – original draft, A.T. and J.J.S.; writing – review & editing, A.T. and J.J.S.; visualization, A.T. and J.J.S.; supervision, J.J.S.; funding acquisition, J.J.S.

## Data availability

The data supporting this article have been included as part of the electronic supplementary information.

## Acknowledgements

We are grateful for support from the Office of Basic Energy Sciences of the U.S. Department of Energy grant DE-SC0014462 and National Science Foundation grant 2223678. Research was also sponsored by the Army Research Office and was accomplished under Grant Number W911NF-22-1-0239. The views and conclusions contained in this document are those of the authors and should not be interpreted as representing the official policies, either expressed or implied, of the Army Research Office or the U.S. Government. The U.S. Government is authorized to reproduce and distribute reprints for Government purposes notwithstanding any copyright notation herein. *E. coli* EW11 was a gift from Pam Silver (Harvard University). Finally, thanks to George N. Bennett for stimulating discussions.

## References

1 F. L. Sousa, T. Thiergart, G. Landan, S. Nelson-Sathi, I. A. C. Pereira, J. F. Allen, N. Lane and W. F. Martin, Phil. Trans. R. Soc. B, 2013, 368, 20130088.

2 I. J. Campbell, G. N. Bennett and J. J. Silberg, Front. Energy Res., 2019, 7, 79.

3 J. T. Atkinson, I. Campbell, G. N. Bennett and J. J. Silberg, Biochemistry, 2016, 55, 7047–7064.

4 K. Syed, CIMB, 2024, 46, 9659–9673.

5 A. M. Terauchi, S.-F. Lu, M. Zaffagnini, S. Tappa, M. Hirasawa, J. N. Tripathy, D. B. Knaff, P. J. Farmer, S. D. Lemaire, T. Hase and S. S. Merchant, Journal of Biological Chemistry, 2009, 284, 25867–25878.

6 E. A. Peden, M. Boehm, D. W. Mulder, R. Davis, W. M. Old, P. W. King, M. L. Ghirardi and A. Dubini, Journal of Biological Chemistry, 2013, 288, 35192–35209.

7 C. Cassier-Chauvat and F. Chauvat, Life, 2014, 4, 666–680.

8 B. W. Burkhart, H. P. Febvre and T. J. Santangelo, mBio, 2019, 10, e02807–18.

9 G. Battistuzzi, M. D’Onofrio, M. Borsari, M. Sola, A. L. Macedo, J. J. G. Moura and P. Rodrigues, J. Biol. Inorg. Chem., 2000, 5, 748–760.

10 P. J. Stephens, D. R. Jollie and A. Warshel, Chem. Rev., 1996, 96, 2491–2514.

11 K. N. McGuinness, N. Fehon, R. Feehan, M. Miller, A. C. Mutter, L. A. Rybak, J. Nam, J. E. AbuSalim, J. T. Atkinson, H. Heidari, N. Losada, J. D. Kim, R. L. Koder, Y. Lu, J. J. Silberg, J. S. G. Slusky, P. G. Falkowski and V. Nanda, Proteins, 2024, 92, 52–59.

12 K.-S. Yoon, M. Ishii, T. Kodama and Y. Igarashi, Archives of Microbiology, 1997, 167, 275–279.

13 J. T. Wan and J. T. Jarrett, Archives of Biochemistry and Biophysics, 2002, 406, 116–126.

14 P. Gou, G. T. Hanke, Y. Kimata-Ariga, D. M. Standley, A. Kubo, I. Taniguchi, H. Nakamura and T. Hase, Biochemistry, 2006, 45, 14389–14396.

15 M. Yamamoto, H. Arai, M. Ishii and Y. Igarashi, Biochemical and Biophysical Research Communications, 2003, 312, 1297–1302.

16 O. Guerrini, B. Burlat, C. Léger, B. Guigliarelli, P. Soucaille and L. Girbal, Curr Microbiol, 2008, 56, 261–267.

17 J. Jacquot, A. Suzuki, J. Peyre, R. Peyronnet, M. Miginiac-Maslow and P. Gadal, European Journal of Biochemistry, 1988, 174, 629–635.

18 N. M. Lewis, A. Sarne and K. R. Fixen, mBio, 2023, 14, e02881–22.

19 C. J. Delebecque, A. B. Lindner, P. A. Silver and F. A. Aldaye, Science, 2011, 333, 470–474.

20 J. M. Clomburg, J. E. Vick, M. D. Blankschien, M. Rodríguez-Moyá and R. Gonzalez, ACS Synth. Biol., 2012, 1, 541–554.

21 Y. J. Zhou, Y. Hu, Z. Zhu, V. Siewers and J. Nielsen, ACS Synth. Biol., 2018, 7, 584–590.

22 D. Leister, Plant Physiol., 2019, 179, 778–793.

23 D. C. Ducat, G. Sachdeva and P. A. Silver, Proc. Natl. Acad. Sci. U.S.A., 2011, 108, 3941–3946.

24 H. Eilenberg, I. Weiner, O. Ben-Zvi, C. Pundak, A. Marmari, O. Liran, M. S. Wecker, Y. Milrad and I. Yacoby, Biotechnol Biofuels, 2016, 9, 182.

25 S. Rumpel, J. F. Siebel, C. Farès, J. Duan, E. Reijerse, T. Happe, W. Lubitz and M. Winkler, Energy Environ. Sci., 2014, 7, 3296–3301.

26 Y. Uda, H. Miura, Y. Goto, K. Yamamoto, Y. Mii, Y. Kondo, S. Takada and K. Aoki, ACS Chem. Biol., 2020, 15, 2896–2906.

27 S. Pochekailov, R. R. Black, V. P. Chavali, A. Khakhar and G. Seelig, ACS Synth. Biol., 2016, 5, 662–671.

28 J. T. Atkinson, L. Su, X. Zhang, G. N. Bennett, J. J. Silberg and C. M. Ajo-Franklin, Nature, 2022, 611, 548–553.

29 R. V. Eck and M. O. Dayhoff, Science, 1966, 152, 363–366.

30 B. R. Gibney, S. E. Mulholland, F. Rabanal and P. L. Dutton, Proc. Natl. Acad. Sci. U.S.A., 1996, 93, 15041–15046.

31 S. E. Mulholland, B. R. Gibney, F. Rabanal and P. L. Dutton, Biochemistry, 1999, 38, 10442–10448.

32 J. D. Kim, D. H. Pike, A. M. Tyryshkin, G. V. T. Swapna, H. Raanan, G. T. Montelione Nanda and P. G. Falkowski, J. Am. Chem. Soc., 2018, 140, 11210–11213.

33 A. C. Mutter, A. M. Tyryshkin, I. J. Campbell, S. Poudel, G. N. Bennett, J. J. Silberg Nanda and P. G. Falkowski, Proc. Natl. Acad. Sci. U.S.A., 2019, 116, 14557–14562.

34 M. Boehm, M. Alahuhta, D. W. Mulder, E. A. Peden, H. Long, R. Brunecky, V. V. Lunin, P. W. King, M. L. Ghirardi and A. Dubini, Photosynth Res, 2016, 128, 45–57.

35 I. J. Campbell, D. Kahanda, J. T. Atkinson, O. N. Sparks, J. Kim, C.-P. Tseng, R. Verduzco, G. N. Bennett and J. J. Silberg, ACS Synth. Biol., 2020, 9, 3245–3253.

36 B. Wu, J. T. Atkinson, D. Kahanda, G. N. Bennett and J. J. Silberg, AIChE Journal, 2020, 66, e16796.

37 J. T. Atkinson, I. J. Campbell, E. E. Thomas, S. C. Bonitatibus, S. J. Elliott, G. N. Bennett and J. J. Silberg, Nat Chem Biol, 2019, 15, 189–195.

38 P. C. Fridy, Y. Li, S. Keegan, M. K. Thompson, I. Nudelman, J. F. Scheid, M. Oeffinger, M. C. Nussenzweig, D. Fenyö, B. T. Chait and M. P. Rout, Nat Methods, 2014, 11, 1253–1260.

39 C. I. Stains, J. L. Furman, J. R. Porter, S. Rajagopal, Y. Li, R. T. Wyatt and I. Ghosh, ACS Chem. Biol., 2010, 5, 943–952.

40 Y. Ohmuro-Matsuyama and H. Ueda, Anal. Chem., 2018, 90, 3001–3004.

41 H.-J. Chang, P. Mayonove, A. Zavala, A. De Visch, P. Minard, M. Cohen-Gonsaud and J. Bonnet, ACS Synth. Biol., 2018, 7, 166–175.

42 H.-J. Chang, A. Zúñiga, I. Conejero, P. L. Voyvodic, J. Gracy, E. Fajardo-Ruiz, M. Cohen-Gonsaud, G. Cambray, G.-P. Pageaux, M. Meszaros, L. Meunier and J. Bonnet, Nat Commun, 2021, 12, 5216.

43 R. S. Smith, S. G. Harris, R. Phipps and B. Iglewski, J Bacteriol, 2002, 184, 1132–1139.

44 C. Engler, R. Kandzia and S. Marillonnet, PLoS ONE, 2008, 3, e3647.

45 B. Barstow, C. M. Agapakis, P. M. Boyle, G. Grandl, P. A. Silver and E. H. Wintermute, J Biol Eng, 2011, 5, 7.

46 A. Truong, D. Myerscough, I. Campbell, J. Atkinson and J. J. Silberg, Protein Science, 2023, 32, e4746.

47 A. W. Reinke, R. A. Grant and A. E. Keating, J. Am. Chem. Soc., 2010, 132, 6025–6031.

48 S. W. Michnick, M. K. Rosen, T. J. Wandless, M. Karplus and S. L. Schreiber, Science, 1991, 252, 836–839.

49 C. R. Robinson and R. T. Sauer, Proc. Natl. Acad. Sci. U.S.A., 1998, 95, 5929–5934.

50 J. S. Klein, S. Jiang, R. P. Galimidi, J. R. Keeffe and P. J. Bjorkman, Protein Engineering Design and Selection, 2014, 27, 325–330.

51 A. K. Shiau, D. Barstad, P. M. Loria, L. Cheng, P. J. Kushner, D. A. Agard and G. L. Greene, Cell, 1998, 95, 927–937.

52 T. De Meyer, S. Muyldermans and A. Depicker, Trends in Biotechnology, 2014, 32, 263–270.

53 T. H. Segall-Shapiro, P. Q. Nguyen, E. D. Dos Santos, S. Subedi, J. Judd, J. Suh and J. J. Silberg, Journal of Molecular Biology, 2011, 406, 135–148.

54 J. T. Atkinson, A. M. Jones, V. Nanda and J. J. Silberg, Protein Engineering, Design and Selection, 2019, 32, 489–501.

55 P. Q. Nguyen, S. Liu, J. C. Thompson and J. J. Silberg, Protein Engineering Design and Selection, 2008, 21, 303–310.

56 J. T. Atkinson, M. S. Chavez, C. M. Niman and M. Y. El-Naggar, Microbial Biotechnology, 2023, 16, 507–533.

57 I. J. Campbell, J. T. Atkinson, M. D. Carpenter, D. Myerscough, L. Su, C. M. Ajo-Franklin and J. J. Silberg, Biochemistry, 2022, 61, 1337–1350.

