## Supporting Information for "Regulating ferredoxin electron transfer using nanobody and antigen interactions"

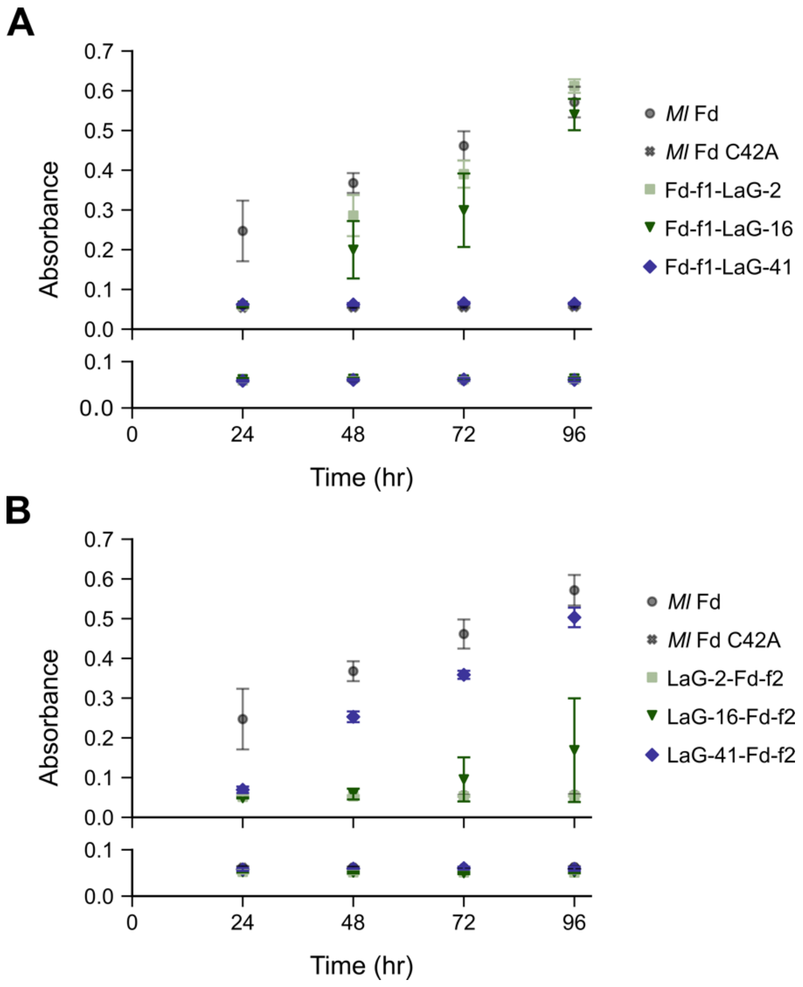


**Figure S1. Growth complementation of *E. coli* EW11 in selective medium across a four-day incubation.** Growth complementation of the Fd split after residue 35 at different times, including (**A**) three Fd-f1-Nb and GFP-Fd-f2 variants and (**B**) three Fd-f1-GFP and Nb-Fd-f2 variants. Data points represent the mean absorbance values of cultures following 24, 48, 72, and 96 hours of growth in m9sa. The top panel represents growth in the presence of inducers while the bottom represent growth in the absence. Data represents the average of three biological replicates, with error bars representing ±1 standard deviation.

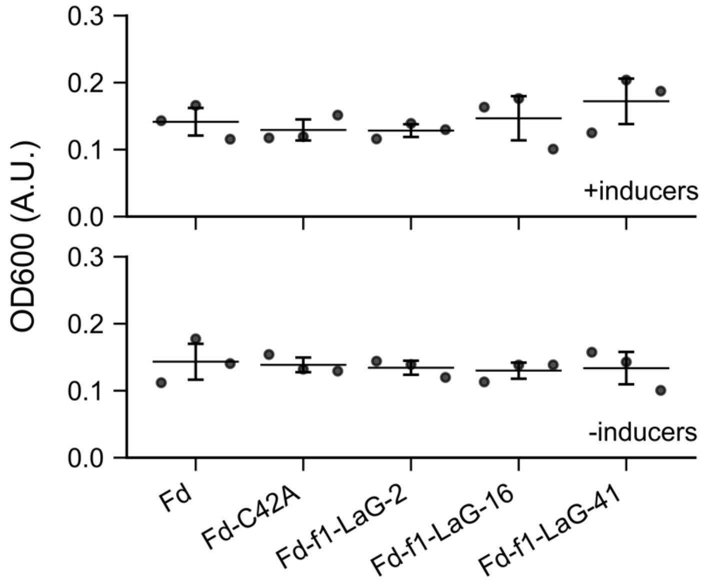


**Figure S2.** ***E. coli*** **EW11 growth in non-selective medium when expressing sFd-35 fragments having Fd-f1 fused to Nbs and Fd-f2 fused to GFP.** *E. coli* EW11 expressing different Fd-f1-Nb and GFP-Fd-f2 variants (split after residue 35) were grown overnight at 37°C in non-selective m9c media which includes sulfur-containing amino acids. No significant differences ($p > 0.1$, two-tailed Welch's t-test) in optical density are observed between cultures in which expression of split ferredoxin fusions from Figure 1C are induced (*top*) or not induced (*bottom*).

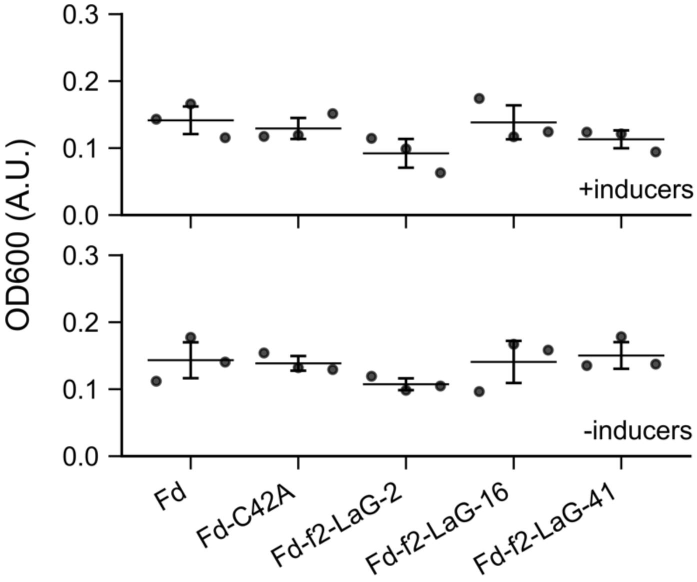


**Figure S3.** ***E. coli*** **EW11 growth in non-selective medium when expressing sFd-35 fragments having Fd-f1 fused to GFP and Fd-f2 fused to Nbs.** *E. coli* EW11 expressing different Fd-f1-GFP and Nb-Fd-f2 variants (split after residue 35) were grown overnight at 37°C in non-selective m9c media which includes sulfur-containing amino acids. No significant differences ($p > 0.1$, two-tailed Welch's t-test) in optical density are seen between cultures in which expression of split ferredoxin fusions from Figure 1D are induced (*top*) or not induced (*bottom*).

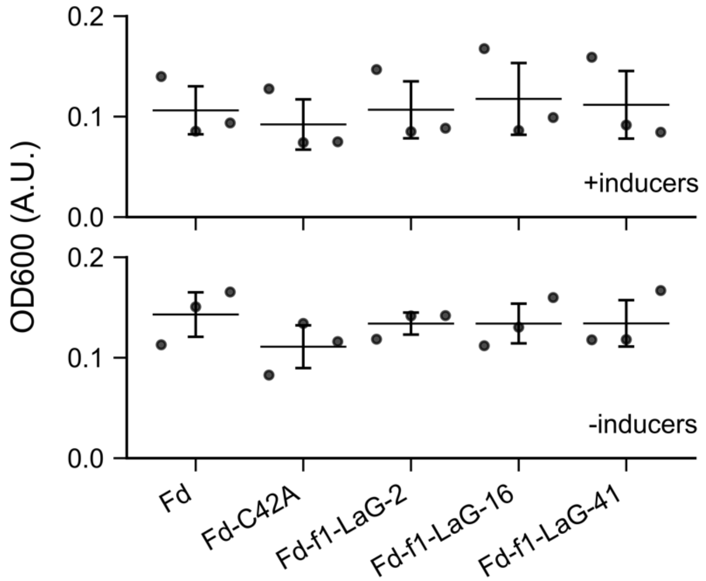


**Figure S4.** ***E. coli*** **EW11 growth in non-selective medium when expressing sFd-55 fragments having Fd-f1 fused to Nbs and Fd-f2 fused to GFP.** Cells expressing different Fd-f1-Nb and GFP-Fd-f2 variants were grown overnight at 37°C in non-selective medium which includes sulfur-containing amino acids. No significant difference in optical density ($p > 0.1$; two-tailed Welch's t-test) are observed between cultures expressing the variants from Figure 2A in the presence (*top*) or absence (*bottom*) of inducers.

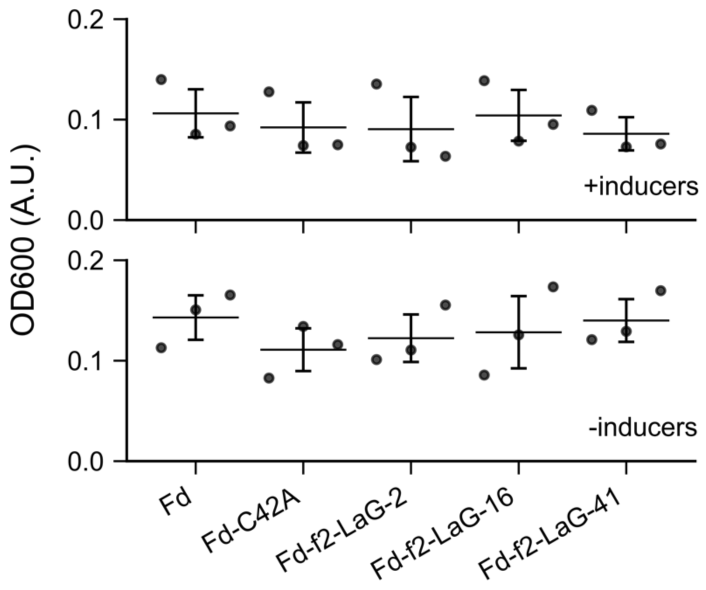


**Figure S5.** ***E. coli*** **EW11 growth in non-selective medium when expressing sFd-55 fragments having Fd-f1 fused to GFP and Fd-f2 fused to Nbs.** Cells expressing different Fd-f1-GFP and Nb-Fd-f2 variants were grown overnight at 37°C in non-selective m9c medium which includes sulfur-containing amino acids. No significant differences in optical density are observed between expressing the split Fd fusions from Figure 2B in the presence (*top*) or absence (*bottom*) of inducers ($p > 0.05$, two-tailed Welch's t-test).

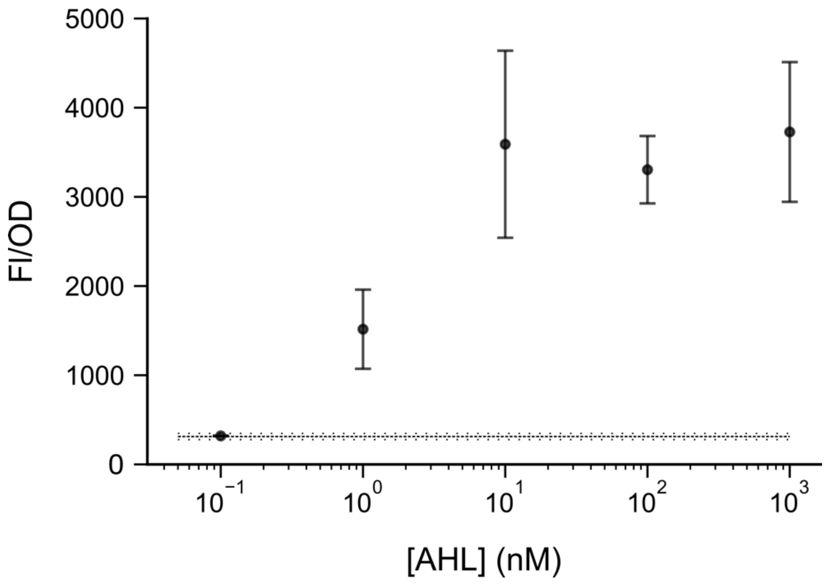


**Fig. S6. GFP induction curve.** *E. coli* EW11 harboring the vector for GFP expression (pAT019) were induced with 0, 0.1, 1, 10, 100, and 1000 nM AHL in non-selective M9c medium. After 20 hours at 37°C, fluorescence (FI) to optical density (OD) were measured and the ratio was calculated. The data represent the mean from three biological replicates, while error bars represent ±1 standard deviation. The line represents the FI/OD ratio for cells grown in the absence of inducer.

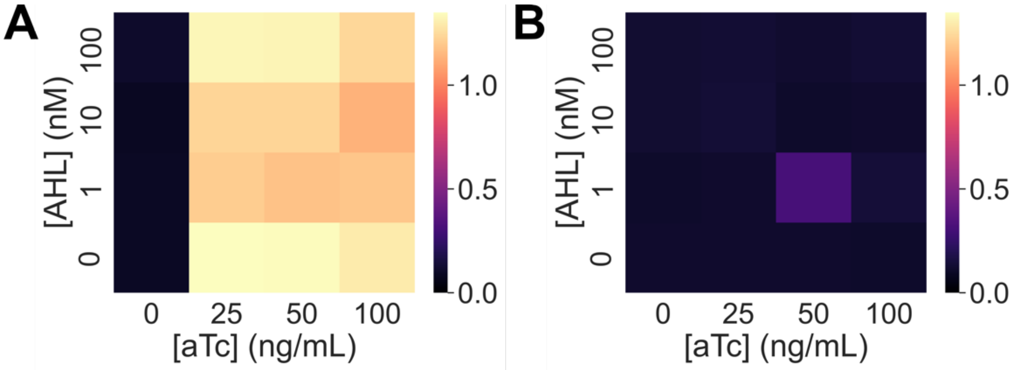


**Fig. S7. Effect of inducing GFP expression on Fd complementation.** GFP was co-expressed with (**A**) native Fd and (**B**) a Fd-C42A mutant. *E. coli* EW11 were incubated at 37°C for 48 hours in selective medium containing varying concentrations of aTc (0, 25, 50, and 100 ng/mL) and AHL (0, 1, 10, and 100 nM), which induce expression of the Fds and GFP, respectively. Growth complementation was not observed with Fd in the absence of inducer, while all other induction conditions presented similar growth complementation. Data represent the mean of three biological replicates.

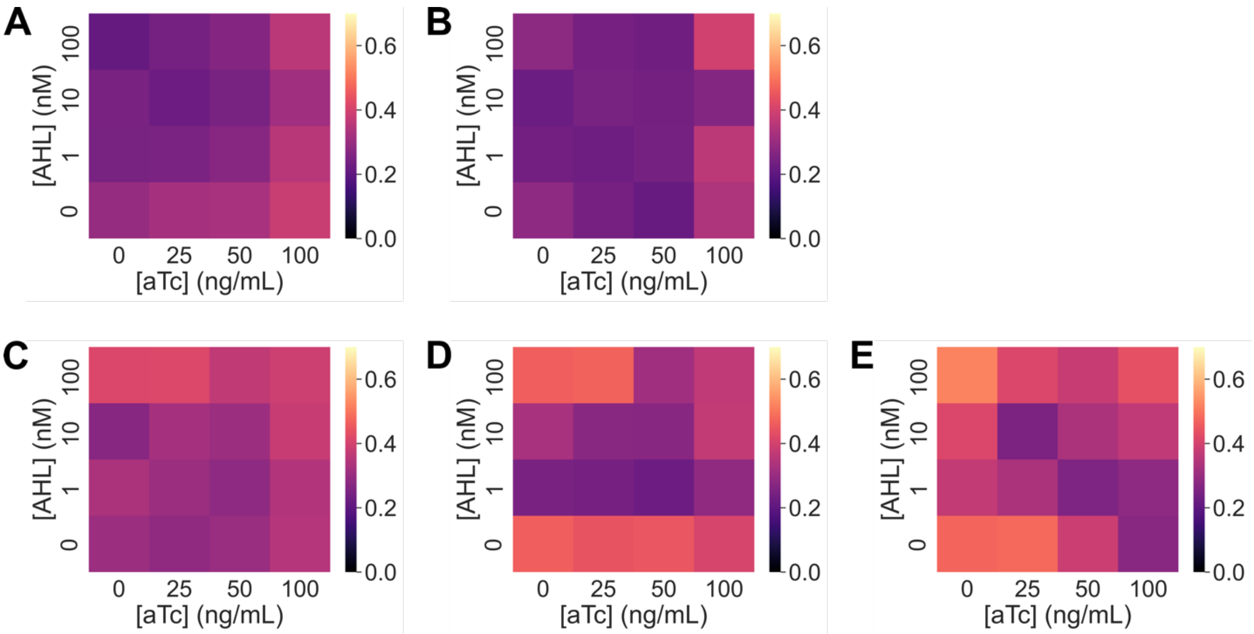


**Fig. S8. Growth of cells expressing domain insertion variants under non-selective conditions.** GFP was co-expressed with (**A**) native Fd, (**B**) Fd-C42A, (**C**) Fd-LaG-2, (**D**) Fd-LaG-16, and (**E**) Fd-LaG-41 in *E. coli* EW11. *E. coli* EW11 were incubated at 37°C for 20 hours in non-selective medium containing varying concentrations of aTc (0, 25, 50, and 100 ng/mL) and AHL (0, 1, 10, and 100 nM), which induce expression of the Fds and GFP, respectively. All experiments were performed using three biological replicates.

**Table S1. Plasmids used in this study.** Listed for each plasmid are the name, antibiotic resistance marker (AbR), origin of replication (ori), promoters (P1 and P2) used to regulate protein expression, and proteins expressed (Protein #1 and #2). For protein fusions, the linker sequences (Linker #1 and #2) are provided. n/a, not applicable.

| Name | AbR | ori | P1 | Protein #1 | Linker #1 | P2 | Protein #2 | Linker #2 |
|---|---|---|---|---|---|---|---|---|
| pFd007 | CmR | ColE1 | Ptet | MI-Fd (1-99) | n/a | n/a | n/a | n/a |
| pFd007-C42A | CmR | ColE1 | Ptet | MI Fd-C42A | n/a | n/a | n/a | n/a |
| pSH001 | KanR | ColE1 | PlasR | GFP mut3b | n/a | n/a | n/a | n/a |
| pSAC01 | SmR | p15A | Pcon | Zea mays FNR | n/a | Pcon | Zea mays SIR | n/a |
| pAT019 | KanR | pSC101 | PlasR | GFP mut3b | n/a | n/a | n/a | n/a |
| pRAP007.35 | CmR | ColE1 | Ptet | Fd(1-35)::FKBP | $(GGGGS)_2AAA$ | Plac | Fd(36-99)::FRB | $(GGGGS)_2AAA$ |
| pAT020 | CmR | ColE1 | Ptet | Fd(1-35)::LaG-2 | $(GGGGS)_2AAA$ | Plac | Fd(36-99)::GFP | $(GGGGS)_2AAA$ |
| pAT021 | CmR | ColE1 | Ptet | Fd(1-35)::GFP | $(GGGGS)_2AAA$ | Plac | Fd(36-99)::LaG-2 | $(GGGGS)_2AAA$ |
| pAT022 | CmR | ColE1 | Ptet | Fd(1-35)::LaG-16 | $(GGGGS)_2AAA$ | Plac | Fd(36-99)::GFP | $(GGGGS)_2AAA$ |
| pAT023 | CmR | ColE1 | Ptet | Fd(1-35)::GFP | $(GGGGS)_2AAA$ | Plac | Fd(36-99)::LaG-16 | $(GGGGS)_2AAA$ |
| pAT024 | CmR | ColE1 | Ptet | Fd(1-35)::LaG-41 | $(GGGGS)_2AAA$ | Plac | Fd(36-99)::GFP | $(GGGGS)_2AAA$ |
| pAT025 | CmR | ColE1 | Ptet | Fd(1-35)::GFP | $(GGGGS)_2AAA$ | Plac | Fd(36-99)::LaG-41 | $(GGGGS)_2AAA$ |
| pAT026 | CmR | ColE1 | Ptet | Fd(1-55)::LaG-2 | $(GGGGS)_2AAA$ | Plac | Fd(56-99)::GFP | $(GGGGS)_2AAA$ |
| pAT027 | CmR | ColE1 | Ptet | Fd(1-55)::GFP | $(GGGGS)_2AAA$ | Plac | Fd(56-99)::LaG-2 | $(GGGGS)_2AAA$ |
| pAT028 | CmR | ColE1 | Ptet | Fd(1-55)::LaG-16 | $(GGGGS)_2AAA$ | Plac | Fd(56-99)::GFP | $(GGGGS)_2AAA$ |
| pAT029 | CmR | ColE1 | Ptet | Fd(1-55)::GFP | $(GGGGS)_2AAA$ | Plac | Fd(56-99)::LaG-16 | $(GGGGS)_2AAA$ |
| pAT030 | CmR | ColE1 | Ptet | Fd(1-55)::LaG-41 | $(GGGGS)_2AAA$ | Plac | Fd(56-99)::GFP | $(GGGGS)_2AAA$ |
| pAT031 | CmR | ColE1 | Ptet | Fd(1-55)::GFP | $(GGGGS)_2AAA$ | Plac | Fd(56-99)::LaG-41 | $(GGGGS)_2AAA$ |
| pAT032 | CmR | ColE1 | Ptet | Fd(1-35)::LaG-2 | $(GGGGS)_1AAA$ | Plac | Fd(36-99)::GFP | $(GGGGS)_2AAA$ |
| pAT033 | CmR | ColE1 | Ptet | Fd(1-35)::LaG-2 | $(GGGGS)_2AAA$ | Plac | Fd(36-99)::GFP | $(GGGGS)_1AAA$ |
| pAT034 | CmR | ColE1 | Ptet | Fd(1-35)::LaG-2 | $(GGGGS)_1AAA$ | Plac | Fd(36-99)::GFP | $(GGGGS)_1AAA$ |
| pAT035 | CmR | ColE1 | Ptet | Fd(1-35)::LaG-2 | $(GGGGS)_3AAA$ | Plac | Fd(36-99)::GFP | $(GGGGS)_2AAA$ |
| pAT036 | CmR | ColE1 | Ptet | Fd(1-35)::LaG-2 | $(GGGGS)_2AAA$ | Plac | Fd(36-99)::GFP | $(GGGGS)_3AAA$ |
| pAT037 | CmR | ColE1 | Ptet | Fd(1-35)::LaG-2 | $(GGGGS)_3AAA$ | Plac | Fd(36-99)::GFP | $(GGGGS)_3AAA$ |
| pAT038 | CmR | ColE1 | Ptet | Fd(1-35)::GFP | $(GGGGS)_1AAA$ | Plac | Fd(36-99)::LaG-2 | $(GGGGS)_1AAA$ |
| pAT039 | CmR | ColE1 | Ptet | Fd(1-35)::LaG-16 | $(GGGGS)_3AAA$ | Plac | Fd(36-99)::GFP | $(GGGGS)_3AAA$ |
| pAT040 | CmR | ColE1 | Ptet | LaG-2 inserted Fd | $(GGGGS)_2AAA$ | n/a | n/a | n/a |
| pAT041 | CmR | ColE1 | Ptet | LaG-16 inserted Fd | $(GGGGS)_2AAA$ | n/a | n/a | n/a |
| pAT042 | CmR | ColE1 | Ptet | LaG-41 inserted Fd | $(GGGGS)_2AAA$ | n/a | n/a | n/a |
